# Adaptive loss of tRNA gene expression leads to phage resistance in a marine *Synechococcus* cyanobacterium

**DOI:** 10.1101/2024.04.25.591146

**Authors:** Sophia Zborowsky, Ran Tahan, Debbie Lindell

**Author notes:** Living Systems Institute, University of Exeter, United Kingdom (Sophia Zborowsky).

## Abstract

*Synechococcus* is a significant primary producer in the oceans, coexisting with cyanophages which are important agents of mortality. Bacterial resistance against phage infection is a topic of significant interest, yet little is known for ecologically relevant systems. Here we use exogenous gene expression and gene disruption to investigate mechanisms underlying intracellular resistance of marine *Synechococcu*s WH5701 to the Syn9 cyanophage. Despite possessing restriction-modification and Gabija defense systems, neither contributed to resistance. Instead, resistance was primarily driven by insufficient levels of Leu^TAA^ tRNA, preventing translation of key phage genes in a passive, intracellular mode of resistance. Restoring cellular tRNA expression rendered the cyanobacterium sensitive to infection. We propose an evolutionary scenario whereby changes in cell codon usage, acquisition of tRNAs by the phage and loss of cell and phage tRNA expression resulted in an effective means of resistance, highlighting the dynamic interplay between bacteria and phages in shaping their co-evolutionary trajectories.

## Introduction

Bacterial resistance mechanisms against phage infection have been extensively studied over the last several decades. This interest stems from an understanding that phages play a major role in the ecology and evolution of bacterial populations^1,2^. Recently, interest in bacterial phage resistance has increased due to a rise in antimicrobial resistance, driving a resurgence in phage therapy research, where resistance to phages poses a major hurdle for clinical use^3^. Furthermore, bacterial defense systems, such as restriction-modification (R-M) and CRISPR–Cas systems, have been adapted as important tools for biotechnology. However, experimentation to elucidate molecular mechanisms of resistance in ecologically relevant systems is rare.

Studies of resistance commonly focus on finding genes that confer protection against infection^3–6^. However, phage resistance is not exclusively due to these types of “active” innate and acquired defense systems. Loss of function, or “passive” modes of resistance, can also protect bacteria against phage infection. The most well studied examples of the latter are mutations that lead to changes in the cell wall, such as alterations in receptors^7–11^ or loss of extracellular polymers^12,13^, impacting phage adsorption. Intracellular resistance caused by a loss of function has been less well studied. Phages use host intracellular components for replication and infections can fail when these are artificially deleted^14–16^. Yet, there is little evidence that bacteria have naturally evolved resistance through changes in intracellular components that result in the arrest of phage infection inside the cell^17^.

*Synechococcus* and *Prochlorococcus* are marine picocyanobacteria that contribute significantly to global primary production^18^. They coexist with high numbers of cyanophages in the oceans^19^. In some regions cyanophages can be a substantial cause of mortality, impacting cyanobacterial distribution, primary production and ocean biogeochemistry^20–23^. In other regions low levels of infection ensue^23,24^ and coexistence is likely due to a high degree of cyanobacterial resistance to co-occurring cyanophages^7,19,25^.

Previously, we reported that resistance to generalist cyanophages (those with broad host ranges) often acts intracellularly, after adsorption and entry^17^. Syn9 is a marine T4-like cyanomyovirus, with a 177.3 Kb genome encoding 226 putative proteins and six tRNAs with a GC content of 40.5%^26^. Syn9 is a generalist that infects multiple *Synechococcus* strains^17,19,27^. One such sensitive *Synechococcus* strain is the open ocean, phycoerythrin-containing, *Synechococcus* sp. strain WH8102, belonging to clade III of *Synechococcus* subcluster 5.1^28,29^. This cyanobacterium has a 2.43 Mb genome encoding 2,583 proteins with a GC content of 59.4%^30^. It has no currently recognizable defense systems against phages^17^, and is sensitive to many cyanophages^19,31,17^. Syn9 cannot replicate on a suite of other cyanobacteria, including the estuarine, phycoerythrin-lacking, *Synechococcus* sp. strain WH5701, which belongs to *Synechococcus* subcluster 5.2^28,29^. This cyanobacterium has a 2.86 Mb genome encoding 3,129 proteins with a GC content of 66%^30^. This cyanobacterium is resistant to all phages tested on it^17,19^. Its genome contains two known resistance mechanisms, a type I R-M and a Gabija system^5^. Neither system has been tested for its ability to provide resistance to phage infection in *Synechococcus* WH5701.

In this study, our primary objective was to unravel the underlying mechanism of intracellular resistance in *Synechococcus* sp. strain WH5701 against Syn9 infection. We found that the R-M and Gabija active defense systems were ineffective in conferring resistance against Syn9. Rather, resistance stemmed from insufficient levels of Leu^TAA^ tRNA due to dramatically reduced production of Leu^TAA^ tRNA from both the cyanobacterial and cyanophage genomes. Our findings underscore the critical role of tRNA availability in phage sensitivity. Furthermore, they reveal a passive form of resistance where adaptive loss of function of an intracellular host component resulted in resistance to phage infection.

## Results

We used two approaches to investigate the mechanism for resistance to Syn9 infection in *Synechococcus* sp. strain WH5701 (*Synechococcus* WH5701 or WH5701-WT-Resistant). First, we examined the functionality of the cyanobacterium’s two known defense systems. Second, we used the stage at which infection is arrested to discern the nature of the mechanism underlying resistance. Our previous observations revealed that Syn9 enters the cell, replicates its DNA, translates many but not all of its proteins, and begins to assemble capsids, with infection stalling at the late stage of DNA packaging^17^. Untranslated proteins include the neck, tail sheath stabilizer, packaging and capsid-maturing protease proteins^17^. Here, we often compare infection of Syn9 in WH5701-WT-Resistant and its derivatives to that of the sensitive *Synechococcus* sp. strain WH8102 (*Synechococcus* WH8102 or WH8102-WT-Sensitive).

### The role of R-M and Gabija systems

We began by examining the two known defense systems present in *Synechococcus* WH5701. Syn9 DNA degradation in this resistant cyanobacterium occurs many hours after the anticipated end of infection based on sensitive strains^17^, different to the rapid degradation expected for DNA targeting systems like R-M and Gabija^4,32^. Nonetheless, we assessed if they are effective against cyanophages. We inactivated an essential gene within each system in two separate mutants; *hsdR* encoding the predicted nuclease in the R-M system^33^, and *gajB* encoding the predicted helicase in the Gabija system^32^ (Fig. 1a). If either system were responsible for conferring resistance, gene inactivation would lead to phage sensitivity.

**Fig. 1.**
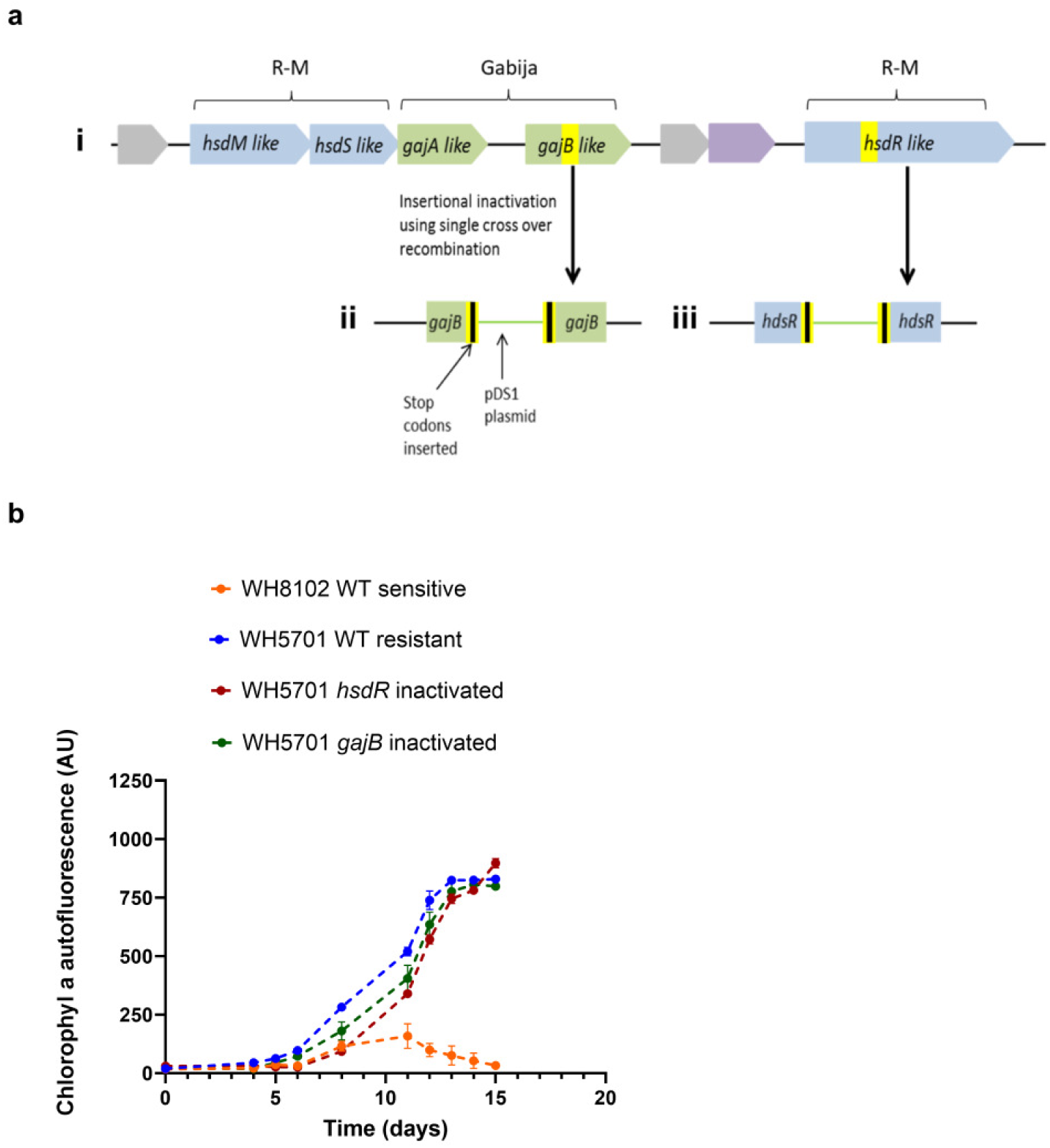
The effect of known defense system genes on the sensitivity of *Synechococcus* WH5701 to the Syn9 phage. **a** Schematic representation of the organization of the two defense systems in the *Synechococcus* WH5701 genome (i) and in the relevant region in the Gabija system (ii) and R-M system (iii) in the mutants after their insertional inactivation with the pDS1 plasmid (see Methods). **b**. Growth of cyanobacteria in the presence Syn9. Average and standard deviation of 5 biological replicates. In (a), blue shows the R-M type I system, green shows the Gabija system, yellow indicates the homologous regions cloned into pDS1, and black indicates where stop codons were inserted. For the R-M system, *hsdM* encodes the modification subunit, *hsdS* the specificity subunit and *hsdR* the endonuclease. For the Gabija system, *gajA* encodes the endonuclease and *gajB* the helicase. The *gajB* and *hsdR* genes were inactivated. AU=arbitrary units.

Upon challenging *Synechococcus* WH5701 inactivation mutants with Syn9 we found no discernible decline in cyanobacterial growth (Fig. 1b). Furthermore, no phage plaques were formed. We next challenged these mutants with two other T4-like cyanophages that the wild-type (WT) cyanobacterium is resistant to but that can attach and enter the cell^17^, S-TIM4 and P-TIM40. Growth of the mutants was not impacted by S-TIM4 (Extended Data Fig.1a) indicating that the R-M and Gabija systems do not confer resistance to this cyanophage either. Growth of the mutants was somewhat reduced by P-TIM40 (Extended Data Fig.1b), suggesting these systems provide some degree of protection against this cyanophage. Consequently, we conclude that, irrespective of whether these two defense systems are active against other cyanophages, neither is responsible for Syn9 resistance. These findings support our previous hypothesis that nuclease-based resistance mechanisms are not responsible for *Synechococcus* WH5701 resistance to Syn9 infection.

### Translational inhibition by tRNA availability

We then asked whether a novel mechanism is involved in phage resistance. Based on the resistance phenotype of Syn9 during infection of *Synechococcus* WH5701, we hypothesized that resistance is due to the absence of crucial proteins required for DNA packaging, tail attachment, and capsid maturation^17^. This could be linked to a failure in their translation, or to their degradation subsequent to translation.

We began by addressing the possibility that the missing proteins are not translated. We hypothesized that a lack of their translation is due to mismatches between codons found in the missing proteins and the cellular tRNA pool. Hence, we compared codon usage between 81 phage proteins detected in *Synechococcus* WH5701 to 35 proteins that were missing (of those detected in WH8102-WT-Sensitive by proteomics)^17^. We found that three codons were significantly more prevalent in non-detected proteins: UUA for Leu^TAA^, AGA for Arg^TCT^, and ACA for Thr^TGT^ (Table 1). Two of these codons, UUA and ACA, are rare in WH5701-WT-Resistant relative to other codons (Table 1, Table S1, Table S2).

**Table 1.**
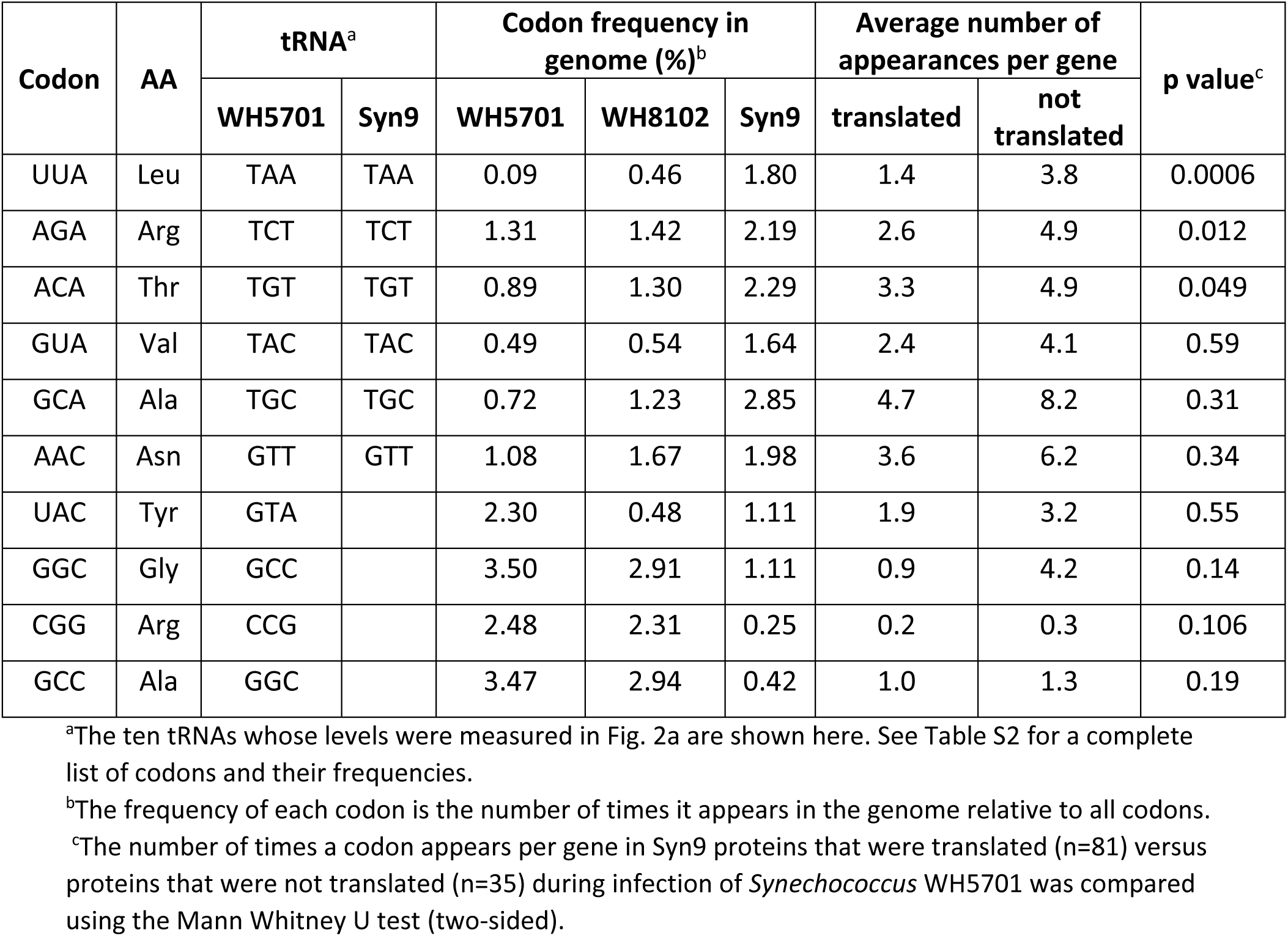
Codon usage and frequency for select tRNAs in *Synechococcus* and Syn9 genomes.

| Codon | AA | tRNA<sup>a</sup> |  | Codon frequency in genome (%)<sup>b</sup> |  |  | Average number of appearances per gene |  | p value<sup>c</sup> |
| --- | --- | --- | --- | --- | --- | --- | --- | --- | --- |
|  |  | WH5701 | Syn9 | WH5701 | WH8102 | Syn9 | translated | not translated |  |
| UUA | Leu | TAA | TAA | 0.09 | 0.46 | 1.80 | 1.4 | 3.8 | 0.0006 |
| AGA | Arg | TCT | TCT | 1.31 | 1.42 | 2.19 | 2.6 | 4.9 | 0.012 |
| ACA | Thr | TGT | TGT | 0.89 | 1.30 | 2.29 | 3.3 | 4.9 | 0.049 |
| GUA | Val | TAC | TAC | 0.49 | 0.54 | 1.64 | 2.4 | 4.1 | 0.59 |
| GCA | Ala | TGC | TGC | 0.72 | 1.23 | 2.85 | 4.7 | 8.2 | 0.31 |
| AAC | Asn | GTT | GTT | 1.08 | 1.67 | 1.98 | 3.6 | 6.2 | 0.34 |
| UAC | Tyr | GTA |  | 2.30 | 0.48 | 1.11 | 1.9 | 3.2 | 0.55 |
| GGC | Gly | GCC |  | 3.50 | 2.91 | 1.11 | 0.9 | 4.2 | 0.14 |
| CGG | Arg | CCG |  | 2.48 | 2.31 | 0.25 | 0.2 | 0.3 | 0.106 |
| GCC | Ala | GGC |  | 3.47 | 2.94 | 0.42 | 1.0 | 1.3 | 0.19 |

<sup>a</sup>The ten tRNAs whose levels were measured in Fig. 2a are shown here. See Table S2 for a complete list of codons and their frequencies.
<sup>b</sup>The frequency of each codon is the number of times it appears in the genome relative to all codons.
<sup>c</sup>The number of times a codon appears per gene in Syn9 proteins that were translated (n=81) versus proteins that were not translated (n=35) during infection of *Synechococcus* WH5701 was compared using the Mann Whitney U test (two-sided).

Next, we wondered whether the resistant *Synechococcus* WH5701 codes for these three tRNAs. We found genes for all three tRNAs in the genome of this cyanobacterium. We then asked whether insufficient tRNA availability could be preventing translation of the missing phage proteins. We used reverse-transcriptase quantitative PCR (RT-qPCR) to measure tRNAs levels in uninfected WH5701-WT-Resistant and compared them to those in WH8102-WT-Sensitive. Two of the three tRNAs (Leu^TAA^ and Arg^TCT^) were below the limit of detection. This is more than 4 orders of magnitude below tRNA levels for common codons in WH5701-WT-Resistant and for these same tRNAs in WH8102-WT-Sensitive (Fig. 2a). The third tRNA (Thr^TGT^) was found at high levels in WH5701-WT-Resistant despite the corresponding codon being rare in this strain. A fourth tRNA that is relatively rare in WH5701-WT-Resistant, Val^TAC^ (Table S1), was also below detection limits, but its codon is not differentially distributed in present and absent Syn9 proteins (Table 1). Despite the occurrence of the relevant tRNA genes in the genome of *Synechococcus* WH5701, tRNA levels of some of them were undetectable in uninfected cells. Thus, this phenomenon is irrespective of the presence of the cyanophage.

**Fig. 2.**
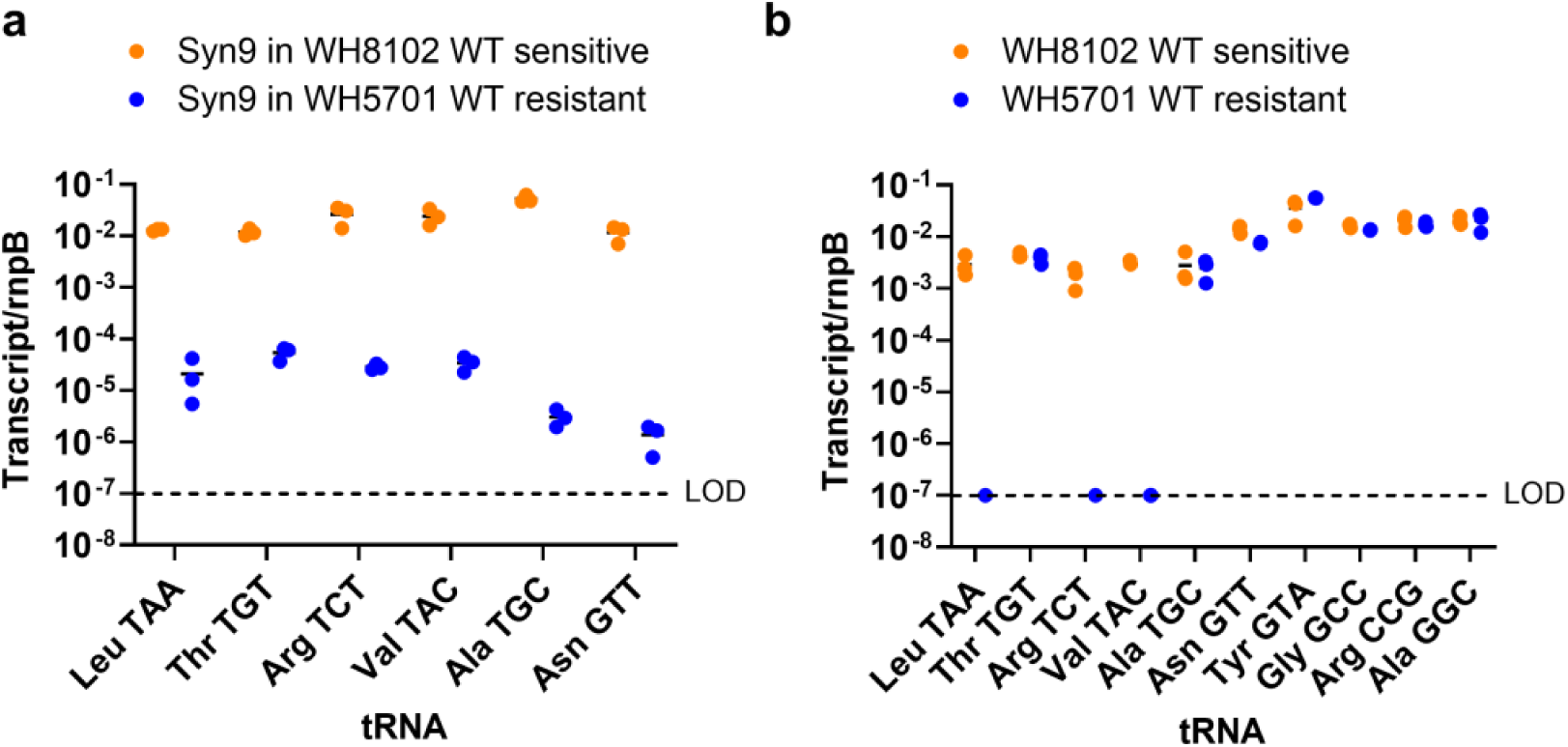
Phage and cyanobacterial tRNA levels. **a.** Levels of cyanobacteria-encoded tRNAs during growth (without infection) for the 6 tRNA genes which have homologues in the Syn9 phage (Leu^TAA^, Thr^TGT^, Arg^TCT^, Val^TAC^, Ala^TGC^, Asn^GTT^) and 4 tRNA genes with common codons in the cyanobacterial strains (Tyr^GTA^, Gly^GCC^, Arg^CCG^ and Ala^GGC^). **b**. Levels of the phage-encoded tRNAs during infection of the resistant *Synechococcus* WH5701 WT and sensitive *Synechococcus* WH8102 WT strains, 5 hours after infection with the Syn9 phage. LOD - limit of detection of the RT-qPCR is approximately 100 transcripts. All data is normalized to levels of cellular *rnpB.* Three biological replicates are shown for each tRNA gene with a black line indicating the average.

Phages often carry tRNA genes, including Syn9, which are thought to compensate for the mismatch between their codon usage and the tRNA pool of the host^26,34–37^ or for depletion of host tRNAs^37,38^. Four of the six tRNA genes carried by Syn9 correspond to codons that are rare (<1 %) or very rare (< 0.5 %)^39^ in WH5701-WT-Resistant (Table S1, Extended Data Fig. 2). This is especially for UAA which accounts for only 0.09 % of WH5701-WT-Resistant codons. These tRNAs match codons common in Syn9^26,35^ (Table S1, Extended Data Fig. 2). Therefore, we wondered whether these phage tRNAs were expressed in the resistant cyanobacterium. We found detectable levels of all six phage tRNAs during infection, but these were approximately 1000-fold lower in WH5701-WT-Resistant relative to WH8102-WT-Sensitive (Fig. 2b). Our findings suggest that these phage tRNAs would not be able to compensate for the low tRNA levels resulting from the cyanobacterial genome.

Given the overall low levels of these tRNAs, we hypothesized that *Synechococcus* WH5701 is resistant to Syn9 because the tRNAs required to translate key phage proteins are not present at sufficient levels. This could be due to insufficient tRNA transcription or rapid degradation after transcription. If there is no, or low, transcription then expressing the tRNAs using appropriate regulatory elements would result in tRNA accumulation in the cells. Conversely, if the tRNAs are targeted for post-transcriptional degradation, then additional transcriptional regulatory elements would not result in their accumulation. To test this, we expressed two undetectable WH5701-WT-Resistant tRNA genes, Leu^TAA^ and Arg^TCT^, together from a replicative plasmid under the control of the cyanobacterial *rnpB* promoter (Fig. 3). Our insert included the tRNA sequences together with their native flanking regions since these can influence tRNA maturation^40,41^. The two tRNAs expressed from the vector were present at levels that were at least 4-orders of magnitude above those in WH5701-WT-Resistant (Fig. 3a,b), indicating that the tRNAs were not targeted for degradation post-transcriptionally in this expression strain, even though they still maintained their native flanks. Thus, we consider a lack of transcription to be the more likely cause for the absence of these tRNAs in WH5701-WT-Resistant than post-transcriptional degradation.

**Fig. 3.**
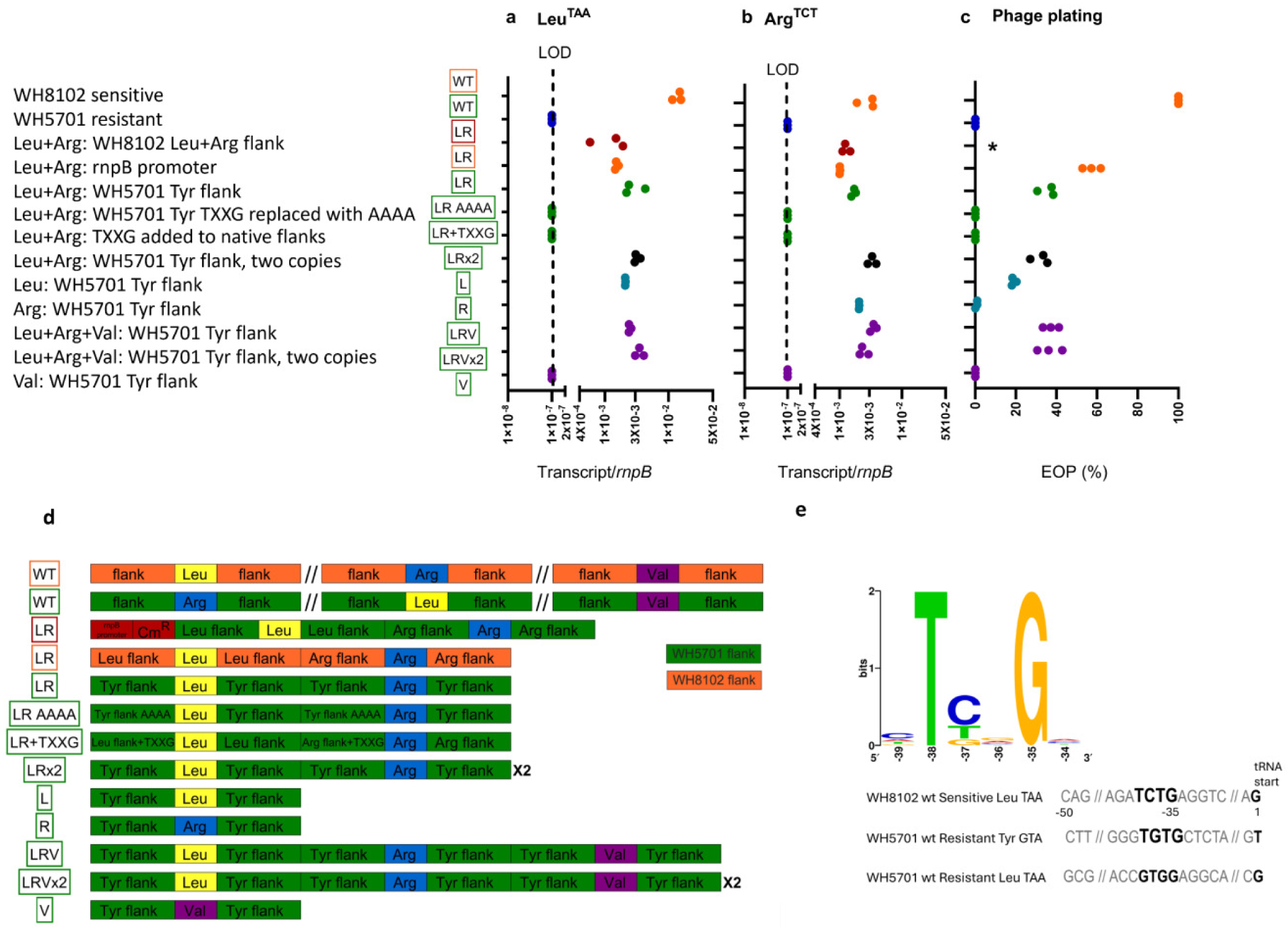
Expression of Leu^TAA^ and Arg^TCT^ tRNAs from, and efficiency of plating on, expression strains. Transcript levels of the Leu^TAA^ tRNA (**a**) and the Arg^TCT^ tRNA (**b**) genes, and efficiency of plating (EOP) of Syn9 on expression strains relative to the sensitive *Synechococcus* WH8102 (**c**). Three biological replicates are shown. Colors match those used in Fig. 4 and 5 and Extended Data Fig. 3, where relevant. LOD, limit of detection (10^-7^ transcripts/*rnpB*). * represents that plaques were obtained, but cannot be quantitatively compared to the rest of the strains. (**d**) A schematic representation of the three tRNA genes relevant for this study, Leu^TAA^, Arg^TCT^, Val^TAC^, and their arrangement and flanking regions in the strains used here. The top two WTs represent the tRNAs in the genomes of *Synechococcus* WH8102-WT-Sensitive (orange) and *Synechococcus* WH5701-WT-Resistant (green), whereas the rest are for the various plasmids introduced into *Synechococcus* WH5701 for construction of the expression strains. The *rnpB* promoter and chloramphenicol resistance gene (Cm^R^) are in red. Flanking regions from the WH8102-WT-Sensitive are in orange, and those from WH5701-WT-Resistant are in green. The tRNA gene the flank was taken from appears inside the flank box. Tyr represents Tyr^GTA^, a tRNA found in high levels in WH5701-WT-Resistant. The tRNA genes on the plasmids were always taken from the WH5701-WT-Resistant strain (**e**) Logo of the TXXG motif and examples of the upstream region of the Leu^TAA^ tRNA gene in the sensitive *Synechococcus* WH8102 and the Tyr^GTA^ tRNA gene in the resistant *Synechococcus* WH5701 where the motif is present, and the same region in the Leu^TAA^ of the latter strain where the motif is missing.

To test if the presence of these tRNAs was sufficient to allow infection, we exposed the expression strain to Syn9. We observed a clear decline in growth of the expression strain, similar to that for WH8102-WT-Sensitive and different to WH5701-WT-Resistant (Fig. 4). Plaques were also formed on lawns of the expression strain (Fig. 4c). These findings unequivocally show that expression of these two tRNAs resulted in phage sensitivity and that *Synechococcus* WH5701 resistance to Syn9 is due, at least partially, to insufficient levels of specific tRNAs. Since this resulted from transcription of the Leu^TAA^ and Arg^TCT^ tRNAs from the cyanobacterial *rnpB* promoter with no other changes to the tRNA genes nor their flanking regions, these findings suggest that neither specific protein degradation nor specific tRNA degradation are the mechanism of resistance.

**Fig. 4.**
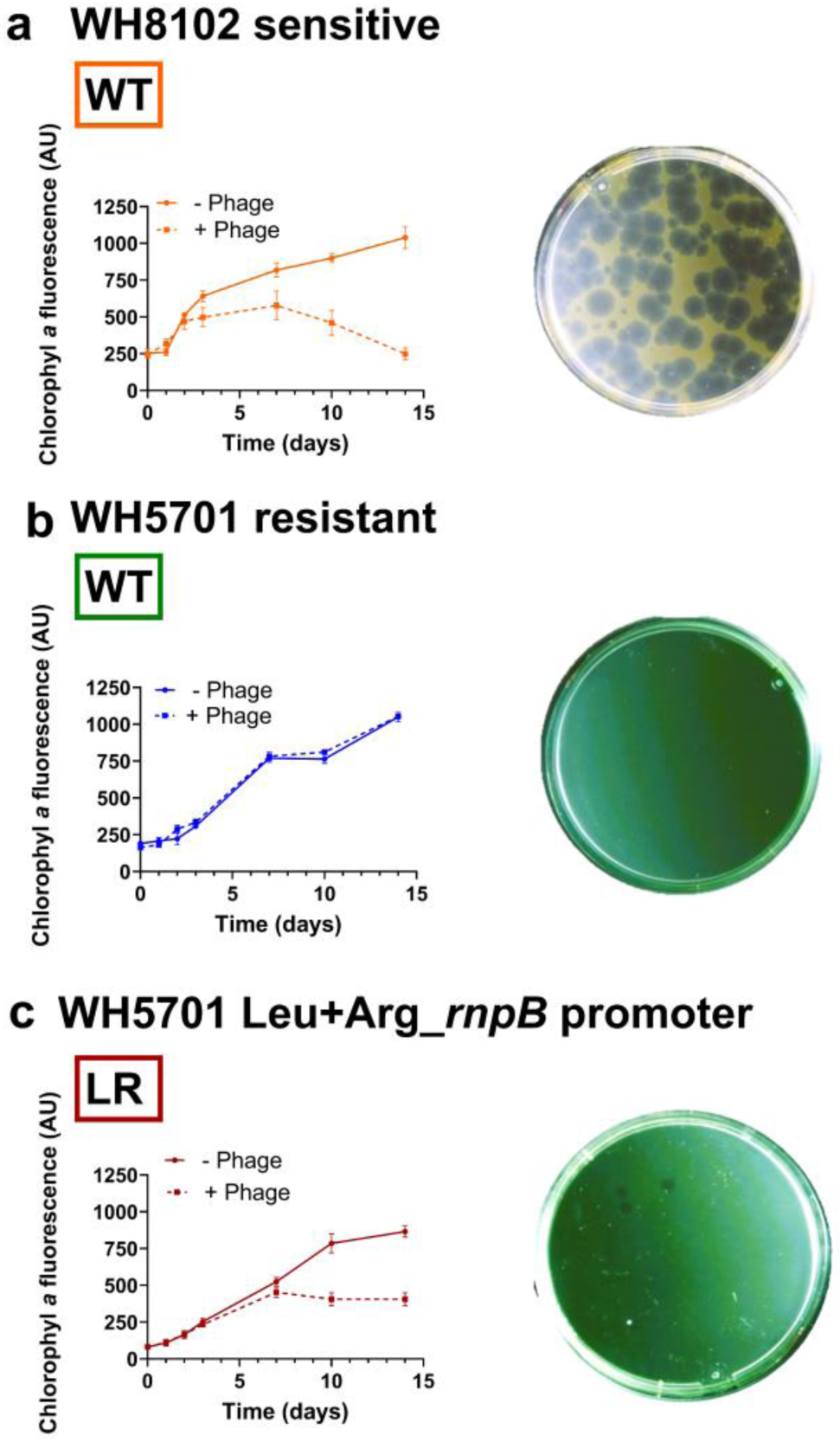
Sensitivity of *Synechococcus* WH5701 tRNA expression strain with *rnpB* promoter to infection by Syn9. Growth of cyanobacterial strains in the presence and absence of Syn9 (left) and plaque formation of Syn9 (right) on sensitive *Synechococcus* WH8102 WT (**a**), resistant *Synechococcus* WH5701 WT (**b**), and WH5701 expression strain where the tRNA genes were cloned under the *rnpB* promoter (**c**). Growth in liquid is average and standard deviation of 5 biological replicates. Plates with Syn9 plaques on cyanobacterial lawns are representative images of 3 biological replicates. WH8102 and WH5701 refer to WT *Synechococcus* WH8102 and WT *Synechococcus* WH5701, respectively. Boxed color-coded abbreviations (WT and LR) match those used in Fig. 3d showing a schematic representation of the arrangement of elements in the *Synechococcus* WT genomes and in the tRNA expression strain with the *rnpB* promoter.

### *Transcriptional loss of tRNAs in* Synechococcus *WH5701*

We set out to understand what causes the lack of expression of the cyanobacterial tRNAs. We hypothesized that these genes have lost necessary regulatory motifs for their production. To assess this, we cloned the two tRNA genes, Leu^TAA^ and Arg^TCT^, with new flanking regions, replacing them with those of the same tRNA genes from WH8102-WT-Sensitive. We also replaced the flanking regions with those of a tRNA gene with high transcript levels from WH5701-WT-Resistant, Tyr^GTA^ (see tRNA levels in Fig. 2a). These inserts were not cloned with an external promoter to allow assessment of whether the flanking regions themselves contained necessary information for tRNA production (see Fig. 3d for construct overview). Both tRNAs were present in high amounts in both *Synechococcus* WH5701 expression strains (Fig. 3a,b). Thus, these flanking regions were sufficient for high tRNA production.

Next, we looked for motifs upstream of tRNA genes that could be responsible for production. This was done for seven tRNA genes in WH5701-WT-Resistant and ten tRNA genes in WH8102-WT-Sensitive for which high transcript levels were experimentally verified (Fig. 2a). We found a conserved motif, TXXG, 35 nt upstream of the first nucleotide in 14 of the 17 genes coding for transcribed tRNAs (Fig. 3e). This motif was absent from the same position for the three genes of the undetected tRNAs, Leu^TAA^, Arg^TCT^, and Val^TAC^ in WH5701-WT-Resistant (Fig. 3e). To test if this motif was needed for tRNA production, we replaced it with AAAA nucleotides in the upstream region of the inserts with the Tyr^GTA^ gene flanking regions. Indeed, the tRNAs were not detected when TXXG was removed (Fig. 3a,b). However, tRNA levels did not increase when we added this motif to the native Leu^TAA^ and Arg^TCT^ tRNA genes in the appropriate position (Fig. 3a,b). These findings indicate that the TXXG motif is needed but not sufficient for production of these two tRNAs. They further suggest that other regulatory elements in the tRNA flanking regions are necessary for tRNA production, although we did not find other conserved motifs.

We then assessed whether tRNA production with and without this motif affected sensitivity to Syn9. We challenged the various *Synechococcus* WH5701 expression strains with Syn9 and found that high tRNA levels resulted in sensitivity to Syn9 (Fig. 3a,b,c), with plaques being produced on lawns as well as a clear decline in cyanobacterial growth (Fig. 5a,b). For the strains in which tRNA transcripts were not observed (Fig. 3a,b), no sensitivity to the phage was found, with neither a decline in cyanobacterial growth nor plaque formation (Fig. 3c, Fig. 5c,d). These findings indicate that tRNA production is sufficient to render *Synechococcus* WH5701 sensitive to Syn9 infection. However, these same WH5701 tRNA expression strains did not become sensitive to infection by the S-TIM4 or P-TIM40 phages, indicating that this *Synechococcus* strain has additional means of resistance against other cyanophages.

**Fig. 5.**
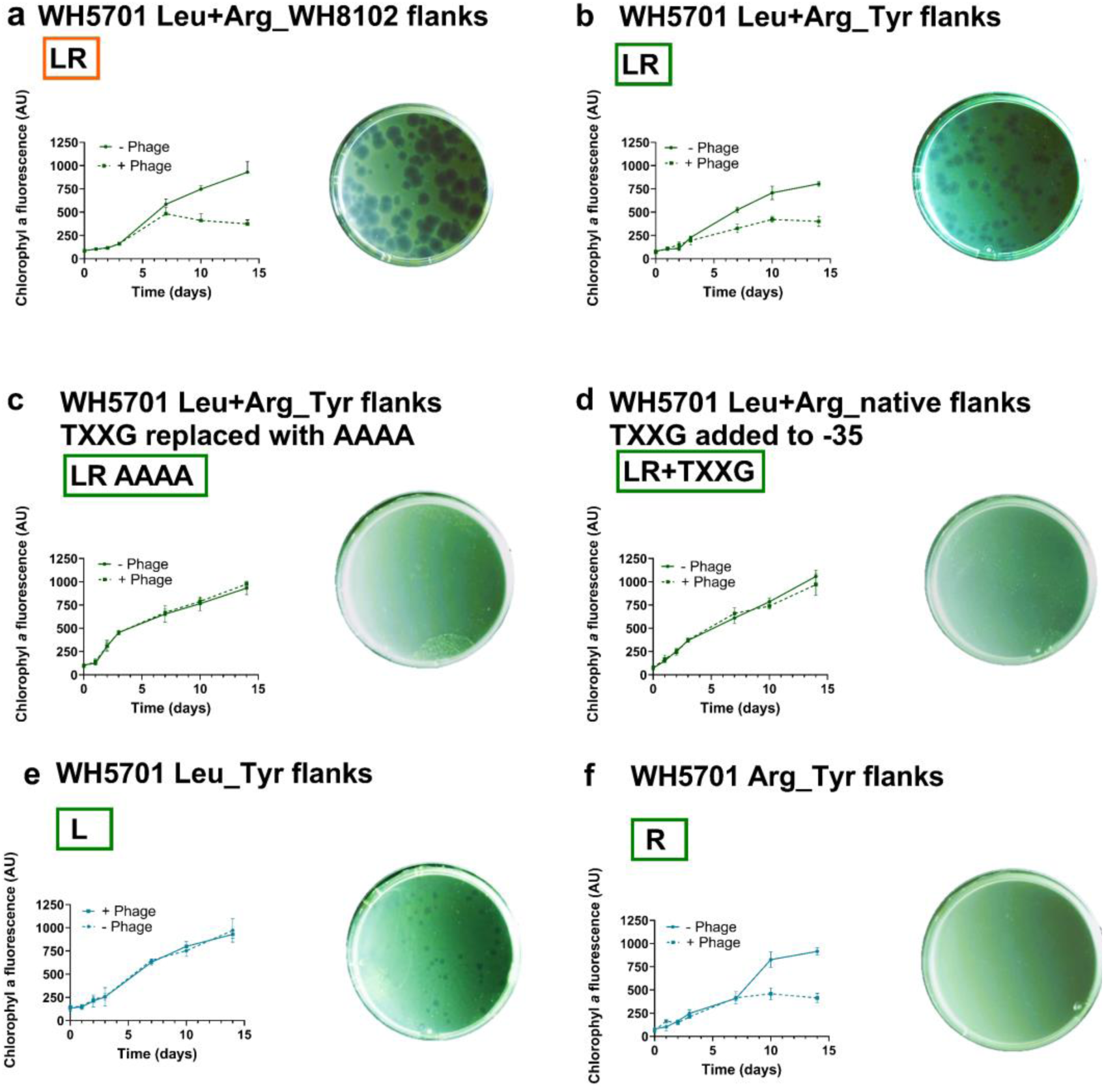
Relevance of tRNA flanking regions for tRNA transcription and phage sensitivity. Growth of cyanobacterial strains in the presence and absence of Syn9 (left) and Syn9 plaque formation (right) on *Synechococcus* WH5701 expression strains (**a**) with the Leu^TAA^ and Arg^TCT^ tRNA genes were cloned with the flanking regions of Leu^TAA^ and Arg^TCT^ of *Synechococcus* WH8102, (**b**), Leu^TAA^ and Arg^TCT^ tRNA genes with Tyr^GTA^ tRNA flanking regions of *Synechococcus* WH5701 (that naturally contain the TXXG motif at the -35 position relative to the first nucleotide of the tRNA), (**c**) Leu^TAA^ and Arg^TCT^ tRNA genes with the Tyr^GTA^ tRNA flanking regions of *Synechococcus* WH5701 where the TXXG motif was replaced with AAAA, (**d**) Leu^TAA^ and Arg^TCT^ tRNA genes with native flanking regions where the TXXG motif was inserted at the -35 position relative to the first tRNA nucleotide, (**e**) with the Leu^TAA^ tRNA gene alone with Tyr^TGA^ flanking regions of *Synechococcus* WH5701, (**f**) with the Arg^TCT^ tRNA gene alone with Tyr^TGA^ flanking regions of *Synechococcus* WH5701. Growth in liquid is average and standard deviation of 5 biological replicates. Plates with Syn9 plaques on cyanobacterial lawns are representative images of 3 biological replicates. WH8102 and WH5701 refer to *Synechococcus* WH8102 and *Synechococcus* WH5701, respectively. Boxed color-coded abbreviations match those used in Fig. 3d for the schematic representation of the arrangement of tRNA genes and flanking regions in the expression strains.

As described above, all three *Synechococcus* WH5701 expression strains with detectable tRNA levels were also sensitive to Syn9. We wondered whether phage production correlates directly to tRNA transcript levels. To assess this, we cloned two copies of the two tRNA genes (Leu^TAA^ and Arg^TCT^), each set with its own Tyr^GTA^ flanking regions, into *Synechococcus* WH5701 (Fig. 3d). We observed a ∼2-fold increase in transcript levels (Fig. 3a,b), but no effect on phage production (Fig. 3c, p=0.7, two tailed U test for EOP). Thus, tRNA transcript levels do not directly correlate with phage production, suggesting other factors are also involved.

We were interested in determining whether both tRNAs were required for the sensitive phenotype. We introduced each tRNA gene separately with the Tyr^GTA^ flanking regions and saw that high levels of Leu^TAA^ alone was sufficient to render *Synechococcus* WH5701 sensitive to Syn9 (Fig. 5e). In contrast, high levels of Arg^TCT^ alone left the cyanobacterium resistant to Syn9 (Fig. 5f). However, phage production was approximately double with both Leu^TAA^ and Arg^TCT^, with an efficiency of plating of 18% with Leu^TAA^ alone versus 35% with both tRNAs (p=0.05, one tailed U test, Fig. 3c). Hence while high levels of Arg^TCT^ alone did not result in phage sensitivity, this tRNA contributed significantly to phage production.

Finally, we assessed whether production of the Val^TAC^ tRNA also renders *Synechococcus* WH5701 sensitive to infection, since it is also below the limit of detection in this cyanobacterium. We cloned the Val^TAC^ gene with Tyr^GTA^ flanking regions from WH5701-WT-Resistant, both alone and together with the Leu^TAA^ and Arg^TCT^ genes (Fig. 3d). The Val^TAC^ gene alone did not result in reduced cyanobacterial growth or phage production (Extended Data Fig. 3a). Its introduction with the other two tRNA genes resulted in sensitivity to Syn9 infection (Extended Data Fig. 3b), which can be ascribed to Leu^TAA^. All three tRNA genes did not increase phage production beyond that with the other two tRNAs without Val^TAC^ (Fig. 3c). Thus, despite the lack of detection of Val^TAC^ in WH5701-WT-Resistant, its introduction did not render *Synechococcus* WH5701 sensitive to infection. Since, the Val^TAC^ codon is not differentially distributed in translated and untranslated Syn9 proteins during infection of WH5701-WT-Resistant (Table 1), these findings are consistent with the conclusion that codon usage discrepancy of select untranslated proteins is behind *Synechococcus* WH5701 resistance to Syn9.

## Discussion

Our findings indicate that lack of production of Leu^TAA^ in *Synechococcus* WH5701 confers it with resistance to Syn9. This was coupled with low transcript levels of the corresponding tRNA carried by the phage. The lack of cellular tRNA transcripts was due, at least partially, to absence of a motif in the flanking region of the tRNA gene, while the mechanism for phage tRNA decline remains unknown. We conclude that scarcity of Leu^TAA^ prevented translation of key Syn9 proteins, resulting in resistance.

The lack of a particular tRNA could be due to targeted degradation^42^, improper maturation^43^, or lack of transcription^44^. Degradation sites are typically found within the sequence of tRNAs themselves^43–47^, while signals for maturation are found in tRNA flanking regions^40,41^. Our finding of a constitutive lack of cellular tRNA transcripts argues against a targeted or stress-induced mechanism of tRNA degradation, as well as against improper maturation since defective tRNAs would likely be present at some level even with constant turnover^41,43^. Furthermore, our finding of a 1000-fold increase in tRNA levels after addition of the *rnpB* promoter upstream of the native tRNA with its native flanking regions, implies that a lack of transcription rather than targeted degradation or defective tRNA maturation is at play. Thus, the most parsimonious explanation for the absence of cyanobacterial Leu^TAA^ and resistance to Syn9 is the lack of tRNA transcription.

Recent research has uncovered a plethora of active innate and acquired defense systems in bacteria^5,6,48,49^. Passive intracellular modes of resistance due to mutations or loss-of-function remain largely unexplored, with known modes of passive resistance typically involving changes to outer cellular components, preventing phage adsorption^6,7–11^. The mode of resistance revealed here can be considered passive since it resulted from the loss of expression of a cellular component required by the phage and does not affect cell growth^17^.

Intracellular passive modes of resistance may be more common than previously appreciated. Many cyanophages can enter cyanobacteria but fail to complete their infection cycle. For example, *Synechococcus* RS9917 (a clade VIII strain from *Synechococcus* subcluster 5.1), may have a similar mechanism to that reported here for WH5701-WT-Resistant. *Synechococcus* RS9917 exhibits early-stage resistance to Syn9, low phage transcription levels^17^, and severe mismatches to Syn9 codons even for early genes (Table S1, Extended Data Fig. 2). Since codon usage discrepancy alone is insufficient to predict resistance (Extended Data Fig. 2), determining whether this is the case for the RS9917-Syn9 interaction requires additional work. In general, further investigation into intracellular passive resistance will enhance our understanding of phage-bacteria interactions and the evolutionary dynamics underlying resistance to phages.

Active defense systems targeting translation have been described previously. Such systems cleave essential host translation factors^50^ or target tRNAs for degradation^42^. These are abortive infection systems resulting in death of the infected cell and provide protection to the population by preventing the spread of the phage^51^. This is in stark contrast to the passive mode of resistance reported here which does not cause the death of the cyanobacterium. Thus, both active, abortive, defense systems and passive, non-abortive, modes of resistance work at the level of translation inhibition.

Phages often encode tRNA genes in their genomes. Recent studies suggest that in some cases this serves to replenish the pool of host tRNAs degraded by active defense systems or depleted due to degradation of the host genome during phage infection^37,38^. Our findings, however, support previous suggestions that cyanophages carry tRNAs to compensate for differences between phage codon usage and the tRNA pool of the host^26,34–37^.

Restoring tRNA expression rendered *Synechococcus* WH5701 sensitive to Syn9, yet phage production remained lower than in the sensitive strain. This could be due to lower expression of the phage’s tRNAs or other Syn9 transcripts in WH5701-WT-Resistant^17^. Alternatively, additional unidentified defense systems could contribute to Syn9 resistance. Likewise, additional defenses could act against other phages, since neither tRNA resistance nor the Gabija or R-M systems provided protection to two other phages. This, together with the fact that no known phage can infect *Synechococcus* WH5701^17,19^ suggests this strain has evolved to possesses multiple layers of defense against cyanophages, culminating in the emergence of a cyanobacterial phage-resistance specialist.

The evolution of loss of tRNA production as a mode of resistance likely occurred gradually. We hypothesize that phages exerted a selective pressure on host codon usage^36,52,53^, resulting in adaptations of cellular codon usage to differ from essential phage replication and morphogenesis genes^34,54,36^. This would have allowed for transcriptional downregulation of key cyanobacterial tRNAs and exerted selective pressure on cyanophages to acquire and express their own tRNA genes^34,36^. We propose that *Synechococcus* WH5701 subsequently evolved an unknown mechanism to limit phage tRNA transcription, rendering phage-acquired tRNAs ineffective and preventing infection. In this multi-step evolutionary scenario, resistance stemmed from the gradual loss of tRNA gene expression, whether encoded by the cell or acquired by the phage, underscoring the dynamic interplay between host and phage in shaping their co-evolutionary trajectories.

## Methods

### Cyanobacterial growth and cyanophage propagation

*Synechococcus* WH5701 and WH8102 cultures were grown in an artificial seawater medium (ASW)^17^. Cultures were grown at 21°C at a light intensity of 15-25 µmol photons·m^-2^·s^-1^ under a 14:10 light:dark regime. Culture growth was measured using chlorophyll *a* autofluorescence (excitation/emission: 440/680 nm) as a proxy for biomass using a Synergy 2 microplate reader (BioTek). Under these conditions, the growth rates of these strains are 2.5 and 1.5 days for *Synechococcus* WH5701 and WH8102 respectively.

For isolation of conjugated colonies, pour plating was used, the cyanobacterial culture was serially diluted and mixed with ultra-pure low-melting point agarose (Invitrogen) at 0.28% in growth medium which was also supplemented with 1 mM sodium sulfite. The helper bacterium, *Alteromonas sp.* strain EZ55, was added to the plates to obtain high plating efficiency^55^.

The Syn9 cyanophage was propagated by infecting a sensitive strain at a low multiplicity of infection (MOI) of <0.01 and allowing the culture to clear. Cells were removed from the phage lysates by filtration through a 0.2 μm pore-sized filter. For small volumes (< 20 ml) Acrodisc syringe filters (PALL) were used; for large volumes Nalgene Rapid Flow 50 mm filter units (Thermo Scientific) were used.

The titer of the lysate (the concentration of infective phages) was determined by plaque assay. The phage lysate was serially diluted and mixed with the sensitive *Synechococcus* strain WH8102 at 2×10^6^ cells per plate in pour plates (as described above, but without the addition of the EZ55 helper bacterium) to form a cyanobacterial lawn on which clearings in the lawn (plaques) could be counted. The plates were incubated at cyanobacterial growth conditions. Plaques were counted until no new plaques appeared. Images of plates with plaques presented here were adjusted for contrast and brightness using the Microsoft Photos application. Adjustments were made across the entire image equally for all images.

### *Insertional inactivation of* Synechococcus *WH5701 genes*

The Type I restriction system was inactivated by disrupting the restriction subunit gene (*hdsR*; protein ID WP_071934184.1), while the Gabija system was inactivated by interrupting the helicase gene (*gajB*; protein ID WP_071934305.1). Gene inactivation was achieved through plasmid integration into the host chromosome via a single homologous recombination event following Brahamsha^56^. A 250 bp homologous segment was modified to include stop codons in all three reading frames to ensure gene inactivation (see Fig. 1a) and was cloned into a pBR322-derived plasmid^56^ (named pDS1)^57^. The chloramphenicol resistance gene in this plasmid was codon optimized for expression in cyanobacteria^58^. The plasmid was electroporated into the S17 *E. coli* donor strain^59^ and colonies were selected on LB plates with 100 μg/ml ampicillin. The plasmid was then mobilized into *Synechococcus* WH5701 by conjugation. Colony formation was performed using pour-plating in the presence of 2 μg/ml chloramphenicol. Successful gene interruption was confirmed by PCR using primers within the plasmid and upstream or downstream of the crossover region in the cyanobacterium (see Table S3 for primer sequences). To ensure full segregation (disruption of all copies of the cyanobacterial chromosome) PCR was performed to verify the absence of wild-type gene copies. Disruption of transcription was verified by RT-qPCR using primers flanking the site of recombination (see Table S3 for primer sequences). Further details on RT and qPCR are provided below.

### Analyzing host and phage codon usage and motif searches

Codon usage in the genomes of *Synechococcus* WH5701, *Synechococcus* WH8102, and Syn9 was analyzed with the frequency of each of the 61 codons counted. Additionally, for phages, codon usage per gene was assessed, comparing the average occurrence of codons between translated and untranslated phage proteins in *Synechococcus* WH5701, as per Zborowsky and Lindell^17^. Codon usage of Syn9 early, middle and late expression clusters was calculated individually for each cluster. Division of Syn9 genes to expression clusters was done according to Doron et al.^27^. The determination of codon usage was carried out using the Countcodon program version 4 with default parameters for bacteria (https://www.kazusa.or.jp/codon/countcodon.html). Statistical comparisons of codon usage per gene were conducted using the Mann-Whitney U test.

Analysis of codon usage preferences and correlation of codon frequencies was done as previously described by Limor-Waisberg et al.^35^ and Enav et al.^36^. Correlation of codon frequencies between Syn9 gene expression clusters and each cyanobacterium was calculated using either the Pearson or Spearman correlation coefficient. If the codon usage of the host and the phage expression cluster was approximately normally distributed the Pearson correlation coefficient was calculated. Otherwise, we used the Spearman correlation coefficient. Normal distribution of the data was assessed using the Lilliefors test for normality.

The TXXG motif was identified after manual inspection of the upstream region of the 10 tRNA genes in *Synechococcus* WH5701 for which tRNA levels were investigated using RT-qPCR. The upstream regions of the 7 expressed tRNAs were compared to the 3 tRNA genes that were not expressed in this cyanobacterium. The motif logo for this region was generated using the WebLogo online tool (https://weblogo.berkeley.edu/logo.cgi)^60^ using the 7 expressed tRNAs from *Synechococcus* WH5701 and the 10 expressed *Synechococcus* WH8102 genes (Fig. 2a).

### *Expression of tRNA genes in* Synechococcus *WH5701*

To explore the impact of the absence of Leu^TAA^ and Arg^TCT^ tRNAs on resistance in *Synechococcus* WH5701, these genes were expressed from a replicative plasmid with different regulatory elements. (1) Native tRNA sequences and flanking regions of the Leu^TAA^ and Arg^TCT^ genes as present in the *Synechococcus* WH5701 genome, with the two tRNA genes cloned downstream of the *rnpB* promoter and a chloramphenicol resistance gene. (2) The same tRNA sequences as in (1) with flanking regions of each gene replaced with the flanking regions of its homolog from the sensitive *Synechococcus* WH8102 in a region of the plasmid without a promoter, and (3) with flanking regions of the Tyr^GTA^ tRNA from *Synechococcus* WH5701, which include the TXXG motif in the -35 position relative to the first tRNA nucleotide in a region of the plasmid without a promoter. (4) The same as in (3) but with the TXXG motif replaced with AAAA. (5) Native tRNA sequences and flanking regions but with the addition of TXXG to the -35 position. (6) The same as (3) but with two copies of each gene. (7 and 8) The same as in (3) but for only one of the tRNA genes (WH5701 Leu^TAA^ alone Tyr^GTA^ flanks or WH5701 Arg^TCT^ alone Tyr^GTA^ flanks). (9) the Val^TAC^ tRNA gene with the Tyr^GTA^ flanks from *Synechococcus* WH5701. (10) Leu^TAA^, Arg^TCT^ and Val^TAC^ with the Tyr^GTA^ flanks from *Synechococcus* WH5701. (11) Same as (10) but two gene copies of each gene. Constructs were synthesized and cloned into a pUC57-Amp plasmid (Genewiz) and were then inserted into the pDS-ProCAT plasmid^58^ using restriction enzymes. The pDS-ProCAT plasmids were then electroporated into *E. coli* S17 and subsequently conjugated into *Synechococcus* WH5701 as described above. Selection of positive colonies was done as described above, except that *E. coli* S17 colonies harboring the plasmid were selected on LB plates supplemented with 750 μg/ml erythromycin (Em).

### Determining transcription levels by reverse transcription

For assessing tRNA levels and the expression of interrupted R-M and Gabija genes, total RNA was extracted from cyanobacterial cells. Samples were taken 5 hours after phage infection, as were non-infected controls, and were centrifuged at 9,000 g for 15 minutes at 4°C. Cells were immediately flash-frozen in liquid nitrogen and stored at -80°C until RNA extraction.

For RNA extraction, cells were thawed on ice and resuspended in a solution containing 10 mM Tris-HCl pH 8, 100 units of RNase inhibitor (Applied Biosystems), and 15,000 units of Lysozyme (Sigma). After incubating at 37°C for 30 minutes for initial cell wall degradation, RNA was isolated using the Quick RNA Mini Prep kit (Zymo). To remove residual DNA, the Turbo DNA Free kit (Invitrogen) was used, incubating with DNase at 37°C for 1 hour. Reverse transcription (RT) was conducted with random hexamer primers using the High Capacity cDNA Reverse Transcription kit (Applied Biosystems). The RT procedure included primer extension at 25°C for 10 minutes, cDNA synthesis at 37°C for 120 minutes, and reaction termination at 85°C for 5 minutes. Samples were diluted five-fold in 10 mM Tris-HCl pH 8 and stored at -20°C before real-time quantitative PCR. The *Synechococcus* WH5701 *rnpB* gene served as a positive control for reverse transcription and as a standardizing marker for gene expression. No RT controls were performed on all samples to ensure that reported transcript levels did not originate from residual phage DNA.

### Real-time quantitative PCR

PCR was conducted using the LightCycler 480 Real-Time PCR System (Roche). The cycling program began with a denaturation step of 95°C for 10 minutes, followed by 35-40 cycles of amplification. Each cycle involved denaturation at 95°C for 10 seconds, annealing at 52°C-68°C for 10 seconds, and elongation at 72°C for 10 seconds. Fluorescence was measured for each reaction (FAM; Ex/Em 465/510nm) at the end of each cycle. The LightCycler 480 software (release 1.5.0) was used to calculate the point at which the fluorescence of a sample exceeded the background fluorescence (Cp) using the absolute quantification/2nd-derivative maximum analysis package. Melting curve analysis on the LightCycler 480 instrument verified the specificity of the amplified PCR product.

Each qPCR reaction contained 1X LightCycler 480 SYBR Green I Master mix (Roche), 200 nM desalted primers, and 5 µl template (in 10 mM Tris-HCl pH8) in a total reaction volume of 20 μl (see Table S3 for primer sequences). The number of DNA copies in the reaction was determined by comparing the Cp to that of a sample with a known DNA concentration using a standard curve. Genomic DNA for standard curves was extracted using the DNeasy Blood & Tissue kit (Qiagen) from cyanobacterial strains and a phenol:chloroform based method for the phages. DNA concentrations in ng/ml were measured by absorbance at 260 nm using a Synergy 2 microplate reader (BioTek) and converted to gene copies/ml by inputting the genome length of the cyanobacterium or phage into the URI Genomics & Sequencing Center calculator for determining the number of copies.

### Determining resistance or sensitivity to phage infection

To assess sensitivity or resistance to each phage, cultures were exposed to phage in liquid medium, and their growth was monitored for 2 weeks. It should be noted that these cyanobacteria have division rates on the order of a doubling every 1.5-2.5 days under the conditions used here and that regrowth consistent with spontaneous mutations that confer resistance appear after approximately a 4-6 weeks in the sensitive *Synechococcus* WH8102 strain. This is well after the period of these experiments.

To maintain consistent phage-bacteria ratios at the beginning of the experiments, cells were quantified using flow cytometry with the Influx flow cytometer (Becton Dickinson), based on forward scatter and autofluorescence (emission at 692/640nm). Yellow-green 1 μm diameter microspheres (Fluoresbrite) served as internal standards for size and fluorescence consistency. Culture growth was monitored using chlorophyll *a* fluorescence (as described above). Furthermore, sensitivity to the phage was tested by plaque assay. A strain was considered sensitive if there was either reduced growth in liquid relative to uninfected controls or visible clearings in plaque assays or deemed resistant if there was no growth reduction in liquid medium and no plaque formation on cyanobacterial lawns.

### Statistical analysis

All statistical analyses were performed using the IBM SPSS statistics version 24. Mann-Whitney’s U test (nonparametric) was used. Normality was tested using the Shapiro Wilk test for normality. Equality of variance was tested using the Levene’s test using SPSS software.

## Data Availability

The data supporting the findings of this study are provided in a combined excel file with separate worksheets for each figure.

## Acknowledgements

We thank Marina Rodnina for the suggestion to investigate codon usage, Gazalah Sabehi, Dror Shitrit, Yoav Arava and Lindell lab members for discussions, and Sarit Avrani and Michael Carlson for comments on the manuscript. This research was supported by funding from the Simons Foundation (SCOPE Grant 329108 and Life Sciences Grant 735081) to D.L. This manuscript is a contribution of the Simons Collaboration on Ocean Processes and Ecology (SCOPE).

## Author contributions

S.Z. and D.L. conceived the project and designed the experiments. S.Z. carried out the experiments, and S.Z. and R.T. analyzed the data. S.Z. and D.L. wrote the manuscript with input from R.T.

---

**Extended Data Fig. 1.**
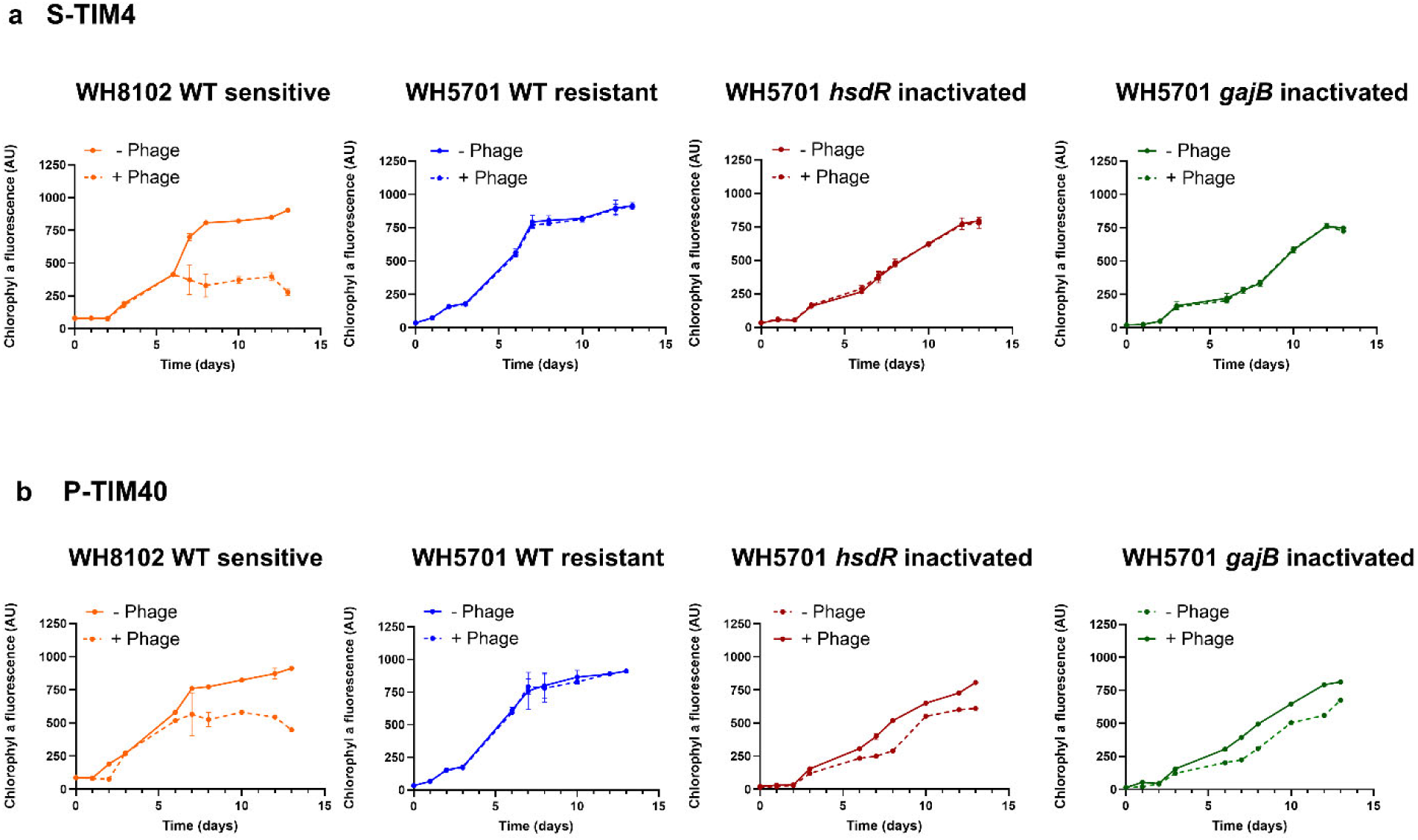
The effect of the R-M and Gabija defense systems on the sensitivity of *Synechococcus* WH5701 to the S-TIM4 and P-TIM40 phages. Growth of cyanobacteria in the presence S-TIM4 (**a**) and P-TIM40 (**b**). Average and standard deviation of 5 biological replicates. For the R-M system, *hsdR* which encodes the endonuclease was inactivated. For the Gabija system, *gajB* which encodes the helicase was inactivated. AU=arbitrary units.

**Extended Data Fig. 2.**
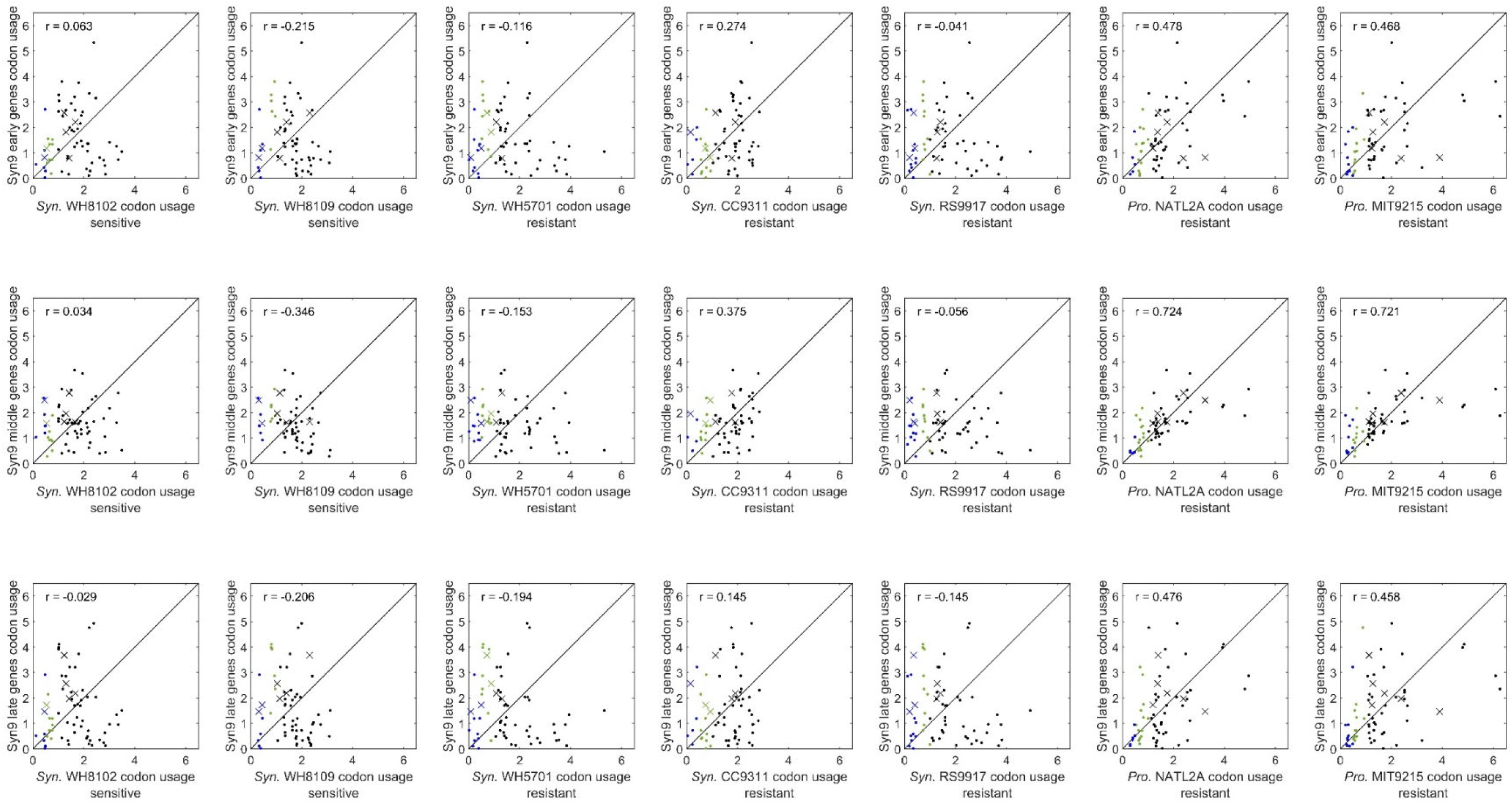
Codon usage preferences of Syn9 gene expression clusters compared to resistant and sensitive cyanobacteria. The frequency of each codon, the number of times it appears in the genome relative to all codons, is shown for sensitive and resistant cyanobacterial strains relative to Syn9 early (top panel), middle (middle panel), and late (bottom panel) transcription clusters. Codons were considered rare if their frequency in the genome is lower than 1% (green) or very rare if their frequency was below 0.5% (blue), following Daniel et al.^39^. Codons for which Syn9 encodes tRNA genes are marked with an X. The cyanobacterial strains are marked as sensitive or resistant following Zborowsky and Lindell^17^. Correlation (r) of codon frequencies between Syn9 gene expression clusters and a cyanobacterium is shown. *Syn.*, *Synechococcus; Pro., Prochlorococcus.* These figures show that there is no direct correspondence between resistance and codon usage correlation as there are sensitive *Synechococcus* strains with no correlation to Syn9 gene clusters and *Prochlorococcus* strains that are resistant to Syn9 with codon usage that correlates with Syn9 gene clusters.

**Extended Data Fig. 3.**
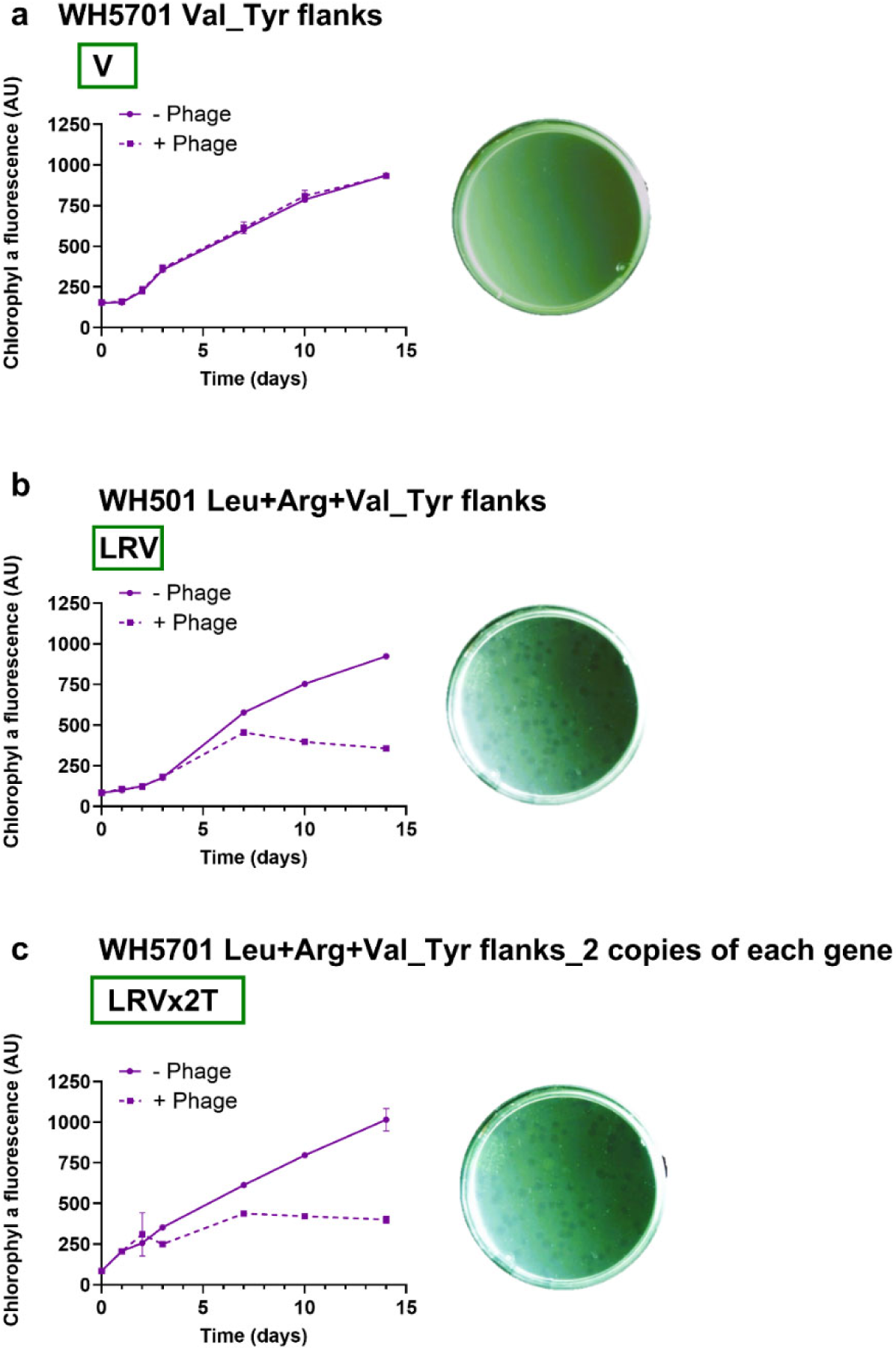
Relevance of tRNA ValTAC for phage sensitivity. Growth of cyanobacterial strains in the presence and absence of Syn9 (left) and Syn9 plaque formation (right) on *Synechococcus* WH5701 expression strains where the ValTAC tRNA gene (**a**), the Leu^TAA^, Arg^TCT^ and ValTAC genes in single copy (**b**) or the Leu^TAA^, Arg^TCT^ and Val^TAC^ genes in two copies (**c**) were cloned with the flanking regions of the WH5701 Tyr^GTA^ tRNA. Growth in liquid is the average and standard deviation of 5 biological replicates. Plates with Syn9 plaques on cyanobacterial lawns are representative images of 3 biological replicates. WH5701 refers to *Synechococcus* WH5701. Boxed color-coded abbreviations match those used in Fig. 3d which shows the schematic representation of the arrangement of tRNA genes and flanking regions in the *Synechococcus* WH5701 plasmid containing strains.

